# Simons Sleep Project (SSP): An open science resource for accelerating scalable digital health research in autism and other psychiatric conditions

**DOI:** 10.1101/2025.02.22.639641

**Authors:** Micha Hacohen, Adam Levi, Hadas Kaiser, LeeAnne Snyder, Alpha Amatya, Brigitta B. Gundersen, John E. Spiro, Ilan Dinstein

**Affiliations:** Cognitive and Brain Sciences Department, Ben Gurion University of the Negev; Simons Foundation; Psychology Department, Ben Gurion University of the Negev

## Abstract

Wearable and nearable devices offer a novel opportunity to measure extensive behavioral and neurophysiological data directly from participants in their home environment. The Simons Sleep Project (SSP) was designed to accelerate research into sleep and daily behaviors in individuals with autism using such techniques. This open-science resource contains raw and processed data from Dreem3 EEG headbands, multi-sensor EmbracePlus smartwatches, and Withings Sleep mats, as well as parent questionnaires and daily sleep diaries. Data were collected successfully for >3600 days/nights from 102 adolescents (10-17 years old) with idiopathic autism and 98 of their non-autistic siblings. Whole-exome sequencing data is also available for all participants and their parents. To demonstrate the utility of this extensive dataset, we first present the breadth of synchronized high-resolution data available across multiple sensors/devices. We then demonstrate that objective sleep measures (e.g., total sleep time) from the three devices are more accurate and reliable than parent reported measures and reveal that sleep onset latency (SOL) was the only objectively defined sleep measure that differed significantly between autistic children and their siblings (of those examined in the study). Moreover, SOL was reliably associated with the severity of multiple behavioral difficulties in all children, regardless of autism diagnosis. These results highlight the importance of measuring sleep directly from participants using objective measures and demonstrate the extensive opportunities afforded by the SSP to further study autism and develop new digital phenotyping techniques for multiple research domains.

## Introduction

The use of wearable/nearable devices with activity, sleep, and other health monitoring applications has grown rapidly over the last decade^1^. These devices offer exciting opportunities for research in many scientific fields because they enable objective and direct quantification of behavior and physiology from large cohorts of participants in ecological settings (i.e., during daily life). The analysis of data from these devices requires the development of automated algorithms that can identify and quantify specific behaviors and physiological events of interest (e.g., tantrums, physical exercise, epileptic seizures, and sleep)^2–5^. To develop accurate, reliable, and transparent algorithms, the scientific community needs access to shared open resources with raw data from simultaneous recordings of multiple devices. Such developments will ultimately be valuable, for example, for performing large longitudinal natural history studies, revealing genotype-phenotype relationships, and developing objective clinical outcome measures.

The goal of the Simons Sleep Project (SSP) is to create a resource for developing such techniques within the context of autism research and an initial focus on studying sleep behavior and neurophysiology. Autism is a behavioral diagnosis that is based on the presence of two core symptoms, social communication difficulties and restricted and repetitive behaviors (RRBs)^6^.

The autism population, however, is highly diverse and most individuals exhibit one or more co-occurring symptoms, which include language delays, intellectual disability, challenging behaviors (e.g., tantrums), sensory problems, hyperactivity, sleep disturbances, and a wide variety of mental health problems (e.g., depression and anxiety)^7,8^. These additional symptoms can be equally or more debilitating for autistic individuals and their families than the core symptoms. Currently available wearable/nearable devices will be particularly useful for quantifying the severity of co-occurring symptoms, studying their etiology and underlying mechanisms, characterizing their relationship with core autism symptoms, and developing new techniques for prevention and intervention. Insights from studying autism should be applicable to a wide variety of neurodevelopmental, psychiatric, and neurological disorders with overlapping symptoms.

Sleep disturbances are one common co-occurring symptom in autism, with previous questionnaire-based studies reporting that >50% of individuals with autism have severe sleep disturbances^9,10^ in contrast to ∼20% of typically developing individuals^11^. The most frequently reported sleep disturbances include difficulties falling asleep, frequent awakening throughout the night, and shorter overall sleep duration^12^. Studies have reported that the severity of sleep disturbances are weakly correlated with the severity of cognitive difficulties and core ASD symptoms^13^ and moderately correlated with sensory problems and challenging behaviors^14–16^. These studies suggest that in some autistic individuals, sleep disturbances may exacerbate the severity of other symptoms, including core symptoms, which may also be generated by shared underlying neurophysiology. In rodents, sleep deprivation during early development has been shown to generate autism-like behaviors^17,18^ that, in some cases, may be ameliorated by improving sleep^19^. In children with autism, treatment with melatonin has been shown to improve sleep disturbances and ameliorate challenging behaviors and additional symptoms^20^. A deeper understanding of specific sleep disturbances and their underlying neurophysiology has the potential to reveal important mechanistic and clinical insights for autism and child development more broadly.

An important caveat of the human studies described above is that they primarily used questionnaires to assess sleep disturbances, thereby basing their conclusions on subjective and potentially biased measures of sleep reported by the participants or their caregivers. To address this, several studies have used overnight polysomnography (PSG) recordings to measure sleep directly and objectively from autistic individuals. During PSG recordings participants are connected to multiple electroencephalogram (EEG), electrooculogram (EOG), and electromyogram (EMG) electrodes that record neural activity, enabling direct identification of sleep and wake periods as well as sleep architecture (i.e. sleep staging) and neurophysiology (e.g. sleep spindles). Most PSG studies have reported that autistic individuals exhibit significantly shorter total sleep time (TST) than controls but do not exhibit differences in wake after sleep onset (WASO)^21–24^. This suggests that some, but not all, parent-reported sleep disturbances are apparent in direct measures of sleep. These studies also reported diverse findings regarding sleep neurophysiology, including less REM sleep^23^, fewer sleep spindles^25,26^, and weaker slow wave activity^24^ in autistic participants relative to controls. While PSG studies are considered the ‘gold standard’ for studying sleep, it is important to consider that all PSG studies described above recorded sleep during a single night while participants were connected to bulky equipment as they slept in a sleep laboratory. Since sleep varies considerably across nights^27^, and autistic individuals find changes in routine particularly aversive^28^, a single PSG recording in a sleep clinic may not faithfully represent their typical sleep behavior at home (i.e., low ecological validity).

To objectively estimate sleep in the home environment, additional studies have used actigraphy devices, which estimate sleep from participant movements. A recent meta-analysis of such studies has concluded that sleep onset latency (SOL) was significantly longer (by 12 minutes), and TST was significantly shorter (by 15 minutes) in recorded autistic individuals relative to controls while WASO did not differ across groups^2^. As with PSG findings, these results suggest that some, but not all parent-reported sleep disturbances are apparent in direct measures of sleep. Note that unlike PSG studies, actigraphy studies estimate sleep/wake periods indirectly from participant movements and several studies have demonstrated that these estimates are inaccurate relative to PSG measures, particularly when quantifying SOL^29^ and WASO^30^. This has motivated the introduction of more modern multi-sensor smartwatches that include photoplethysmography (PPG) sensors for measuring heart rate and heart rate variability in addition to measuring movements with accelerometry. The added information from PPG sensors seems to enable higher accuracy in identifying sleep/wake segments and accurately quantifying WASO^31^, but such devices have not been used in autism research to date. Additional nearable devices such as the Withings sleep mat, which is placed under the mattress and senses pressure changes, can provide information regarding time in bed and participant movements that could further increase the accuracy of sleep measures at home^32^. Such devices have also not been used in autism research to date.

The SSP is intended to rapidly accelerate research in autism and other psychiatric disorders by providing the research community with an open data resource that contains raw and processed data from >3600 nights that were recorded with three devices: Dreem3 EEG headband (*Beacon Inc.*), EmbracePlus smartwatch with accelerometer, PPG, electrodermal activity (EDA), and skin temperature sensors (*Empatica Inc.*), and Withings Sleep Mat (*Withings Inc*.). Corresponding parent-reported data were also collected from a daily sleep diary and baseline questionnaires. The devices were selected for their ability to record and save raw sensor-level data, which is critical for performing transparent and reliable research that is not based on data from proprietary algorithms that are closed-source and can change without notice.

We recorded data from 102 autistic children and 98 of their non-autistic siblings (10-17 years old) who live in the same household, thereby constraining multiple familial factors, which are likely to have a strong impact on the children’s sleep environment and sleep hygiene^33^. Note that siblings of autistic children are known to exhibit a considerably higher prevalence of neurodevelopmental and psychiatric conditions relative to the general population^34^, likely because they share 50% of their sibling’s genes. For example, compared to the general population, siblings have a 3.5-fold higher prevalence of attention deficit hyperactivity disorder (ADHD)^35^, which has also been associated with increased sleep disturbances^36^. The SSP should, therefore, be particularly useful for researchers using dimensional^37^ or transdiagnostic^38^ approaches to disentangle the relationships between specific sleep disturbances and specific neurodevelopmental symptom domains, some of which are shared across siblings and others that are not.

The availability of parallel recordings from multiple devices/sensors as well as inclusion of subjective and objective sleep measures offer a unique opportunity to compare measures of sleep behavior and neurophysiology, assess the agreement between objective and subjective sleep measures, and study sleep disturbances in autistic children and their siblings at a level of detail that has not been possible to date. The goal of this report is to introduce the SSP dataset, demonstrate its potential, and facilitate its utilization by the broad research community.

## Results

Data were successfully recorded from 102 autistic children (86 male) and 98 non-autistic siblings (53 male), between the ages of 10 and 17, from 102 families in the Simons Powering Autism Research for Knowledge (SPARK) cohort (Figure 1). Children were recorded in their home setting over multiple days (EmbracePlus) and nights (all devices) with the final dataset comprising 3,630 nights: 1,687 nights recorded with all three devices, 1,125 nights recorded with two devices, and 818 nights recorded with a single device (8,129 device recordings in total). Corresponding sleep diary data is also available for 2,630 of these nights. Note that the counts above refer to Dreem and Withings night-time recordings with at least 3 hours of sleep and EmbracePlus recordings with at least 6 hours of data per 24-hour period (see Methods).

**Figure 1:**
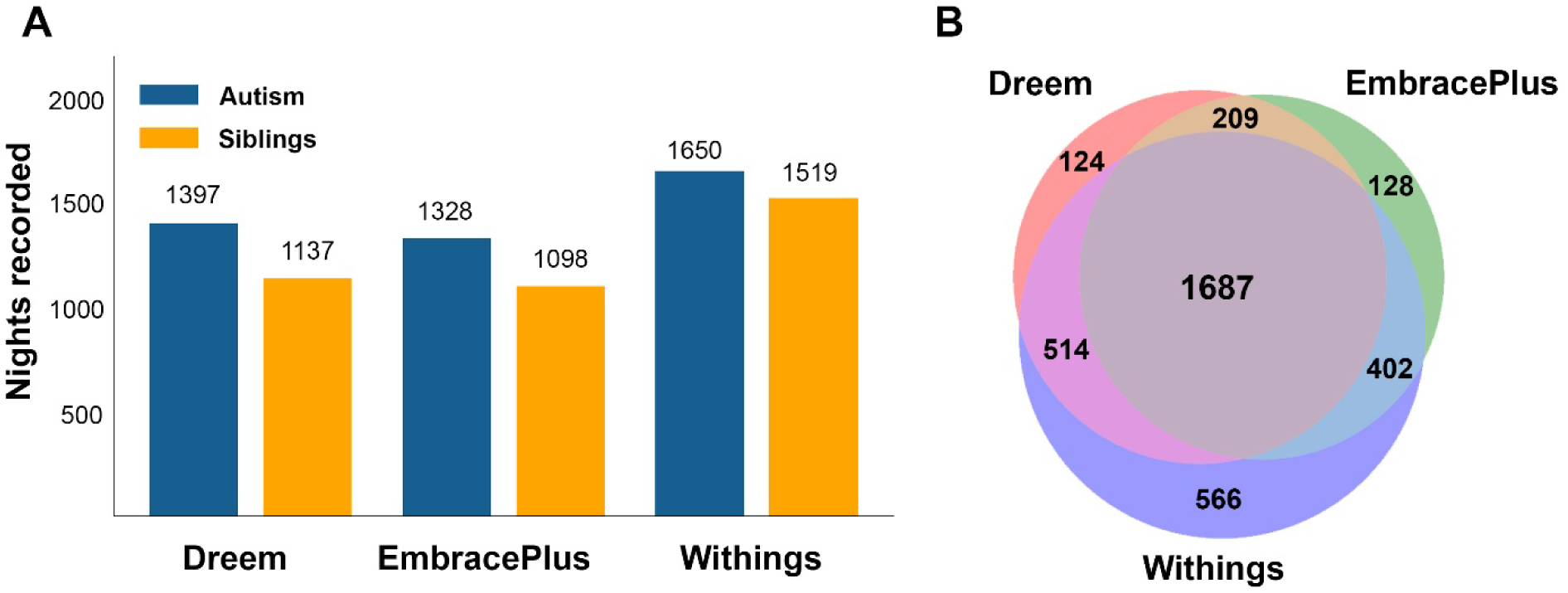
Data overview. **A.** Summary of nights/days recorded with Dreem, EmbracePlus, and Withings devices from the autism (blue) and siblings (yellow) groups. **B.** Venn diagram presenting the number of nights recorded with each device and their overlap (i.e., simultaneous recordings with multiple devices).

### Behavioral characterization of autism and sibling groups

Parents of all participating children completed baseline questionnaires regarding each child’s behavioral abilities and difficulties (Table 1). We examined differences across autism and sibling groups in key measures of core autism symptoms, adaptive behaviors, challenging behaviors, sensory sensitivities, co-occurring psychiatric symptoms, and sleep behaviors (Figure 2). We performed a mixed linear model analysis for each measure to assess differences across autism and sibling groups (fixed effect) while accounting for potential differences in parent reports across families (random effects) as well as the age and sex of participating children (additional fixed effects).

**Figure 2:**
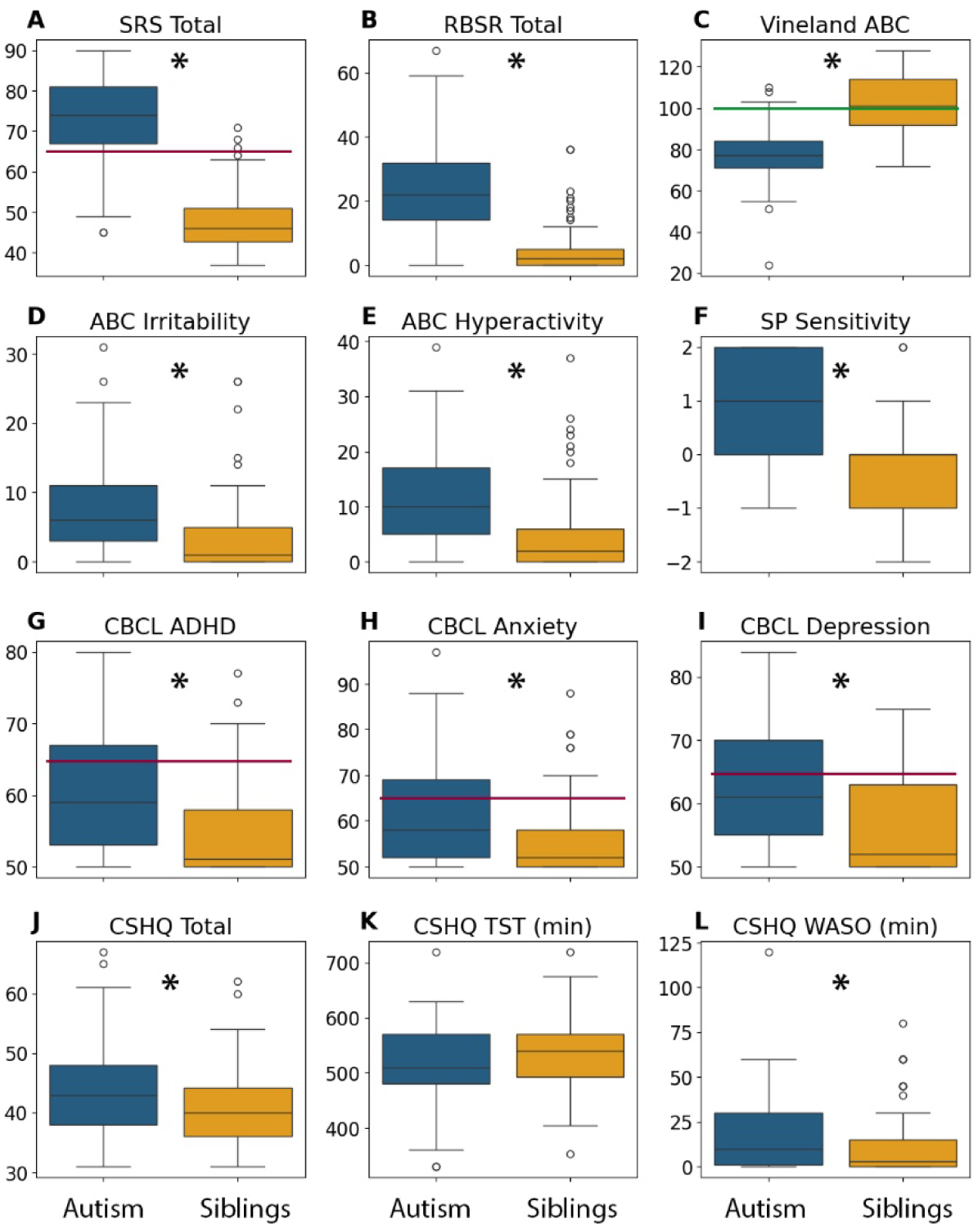
Parent-reported behavioral symptoms and abilities in autistic children (blue) and siblings (yellow). **A.** Social Responsiveness Scale (SRS). **B.** Repetitive Behaviors Scale - Revised (RBS-R) **C.** Vineland Adaptive Behavior Composite (ABC) **D.** Irritability subscale from the Aberrant Behaviors Checklist (ABC) **E.** Hyperactivity subscale from the ABC. **F.** Sensory sensitivity subscale from the Sensory Profile **G.** ADHD symptoms from the Child Behavior Checklist (CBCL) **H.** Anxiety symptoms from the CBCL **I.** Depression symptoms from the CBCL. **J.** Child Sleep Habits Questionnaire (CSHQ) total sleep disturbance score **K.** CSHQ parent-reported TST in minutes **L.** CSHQ parent-reported WASO in minutes. **Red lines:** clinical cutoff (t score = 65). **Green line:** population mean. Asterisks: significant differences between autism and sibling groups according to mixed linear model analyses (p<0.05).

**Table 1:** Participant characteristics.

| Variable | Autism (n=102) | Siblings (n=98) | Statistics |
| --- | --- | --- | --- |
| **Age** | 13.94 ± 2.13 | 13.46 ± 2.16 | t=1.56, p=0.120 |
| **Sex (male/female)** | 86/16 | 52/46 | $\chi^2=21.34$, p=0.000 |
| **SRS-2 total** | 73.71 ± 10.01 | 47.89 ± 7.14 | t=20.83, p=0.000 |
| **RBS-R total** | 24.52 ± 14.05 | 4.15 ± 6.76 | t=12.91, p=0.000 |
| **Vineland** |  |  |  |
| Adaptive behavior composite | 77.64 ± 12.81 | 102.36 ± 14.26 | t=-12.84, p=0.000 |
| Communication | 82.17 ± 14.02 | 104.50 ± 10.55 | t=-12.63, p=0.000 |
| Socialization | 72.15 ± 18.33 | 101.09 ± 12.43 | t=-12.95, p=0.000 |
| Daily Living Skills | 82.05 ± 16.02 | 101.22 ± 16.47 | t=-8.30, p=0.000 |
| **CBCL total** | 52.97 ± 9.82 | 47.31 ± 10.96 | t=3.83, p=0.000 |
| **ABC** |  |  |  |
| Irritability | 8.24 ± 7.29 | 3.46 ± 5.02 | t=5.35, p=0.000 |
| Social withdrawal | 9.42 ± 7.84 | 2.68 ± 4.18 | t=7.50, p=0.000 |
| Stereotypic behavior | 4.39 ± 3.93 | 0.34 ± 1.19 | t=9.74, p=0.000 |
| Hyperactivity | 11.44 ± 7.99 | 4.64 ± 6.85 | t=6.42, p=0.000 |
| Inappropriate speech | 2.84 ± 2.62 | 0.61 ± 1.11 | t=7.74, p=0.000 |
| **CSHQ total** | 46.19 ± 8.05 | 43.31 ± 6.67 | t=2.74, p=0.007 |
| **Sensory Profile 2** |  |  |  |
| Sensory sensitivity | 0.86 ± 0.91 | -0.56 ± 0.94 | t=10.86, p=0.000 |
| Sensory registration | 0.88 ± 0.92 | -0.56 ± 0.99 | t=10.62, p=0.000 |
| Sensory avoidance | 0.91 ± 0.86 | -0.55 ± 0.88 | t=11.79, p=0.000 |
| Sensory seeking | 0.12 ± 0.72 | -0.91 ± 0.83 | t=9.29, p=0.000 |
| **Medical diagnoses** |  |  |  |
| ADHD | 58 (56.86%) | 30 (29.41%) | $\chi^2=12.93$, p=0.000 |
| Depression | 33 (32.35%) | 23 (23.47%) | $\chi^2=1.54$, p=0.21 |
| Anxiety | 16 (15.68%) | 11 (11.22%) | $\chi^2=0.97$, p=0.32 |
| **Medication** |  |  |  |
| Any medication | 66 (69.47%) | 49 (52.69%) | $\chi^2=4.890$, p=0.027 |
| Antidepressants | 31 (32.63%) | 13 (13.98%) | $\chi^2=8.110$, p=0.004 |
| Stimulants | 30 (31.58%) | 13 (13.98%) | $\chi^2=7.285$, p=0.007 |
| Antianxiety | 26 (27.37%) | 13 (13.98%) | $\chi^2=4.343$, p=0.037 |
| Antipsychotics | 12 (12.63%) | 1 (1.08%) | $\chi^2=8.038$, p=0.005 |
| Sleep aids | 8 (8.42%) | 8 (8.6%) | $\chi^2=0$, p=1 |
| Anticonvulsants | 4 (4.21%) | 2 (2.15%) | $\chi^2=0.151$, p=0.698 |

Mean (M), standard deviation (std), number (n), and percentage (%), are reported per measure as relevant. Questionnaires included Social Responsiveness Scale, 2<sup>nd</sup> edition (SRS-2), Repetitive Behaviors Scale – Revised (RBS-R), Vineland adaptive behavior scale, Child Behavioral Check List (CBCL), Aberrant Behaviors Checklist (ABC), Child Sleep Habits Questionnaire (CSHQ), Family Inventory of Sleep Habits (FISH), Sensory Profile, 2<sup>nd</sup> edition.

These analyses revealed significantly higher scores in autistic children relative to their siblings on Social Responsiveness Scale (SRS) total (β = 26.3, p < 0.0001), Repetitive Behaviors Scale - Revised (RBS-R) total (β = 21.1, p < 0.0001), Sensory Profile sensitivity (β = 1.4, p < 0.0001), Aberrant Behaviors Checklist (ABC) irritability ( β = 5.4, p < 0.0001), ABC hyperactivity (β = 6.6, p < 0.0001), Child Behavior Checklist (CBCL) ADHD symptoms (β = 5.33, p < 0.0001), CBCL anxiety symptoms (β = 6.5, p < 0.0001), CBCL depression symptoms (β = 6.27, p < 0.0001), and Child Sleep Habits Questionnaire (CSHQ) total sleep disturbance scores (β = 3.08, p = 0.0018), as well as significantly lower scores on the Vineland adaptive behaviors composite (β = -24.3, p < 0.0001). Parent-reported TST was, on average, 16 minutes shorter in the autistic children relative to their siblings but did not differ significantly across groups (β = -8.42, p = 0.29). Parent-reported WASO was, on average, 8 minutes longer in the autistic children and did differ significantly across groups (β = 7.66, p = 0.006).

There was no significant effect of sex on any of the parent-reported questionnaires or measures (p > 0.1) and a significant effect of age only on ABC irritability (β = -0.05, p = 0.004), ABC hyperactivity (β = -0.04, p = 0.03), and TST (β = -0.56, p = 0.0017). Differences between siblings (within family) were considerably larger than differences across families with Intraclass Correlation Coefficients (ICCs) ranging between 0.16 and 0.37.

### Synchronized recordings with multiple devices

EmbracePlus smartwatch data were recorded continuously when participants wore the device, Dreem3 headband data was recorded when participants activated the device before going to sleep until they turned it off in the morning, and Withings mattress sensor data was recorded automatically whenever participants entered their bed. A single Samsung A51 smartphone was used to manage all devices and synchronized data acquisition to a single clock.

This design enabled us to directly compare data across multiple sensor recordings with high temporal fidelity as demonstrated in a 24-hour recording from a single participant (Figure 3). The devices yielded complementary information regarding the participant’s behavior and sleep/wake state during the night. For example, the activity count time-course, which quantifies movements in 1-minute epochs using the EmbracePlus accelerometer, was correlated with pressure changes recorded by the Withings Sleep mat when the participant was in bed (r =0.41, p < 0.001, Figure 2B). Similarly, sleep and wake epochs identified by the Dreem sleep staging algorithm in the EEG data (Figure 3C) exhibited partial agreement with sleep and wake epochs identified by the EmbracePlus algorithm using accelerometer data (Sensitivity: 0.82, Specificity: 0.27, Figure 3D). Agreement across devices and algorithms varied across participants and nights. Note that the analyses presented here used measures (i.e., activity counts and sleep/wake epoch labels) that were generated by the proprietary algorithms of each device manufacturer. Future studies could utilize the raw data also available in the SSP (e.g., Figure 3E) to quantify activity and identify sleep/wake epochs using alternative algorithms that may yield higher agreement/reliability across devices (also see Figure 6).

**Figure 3:**
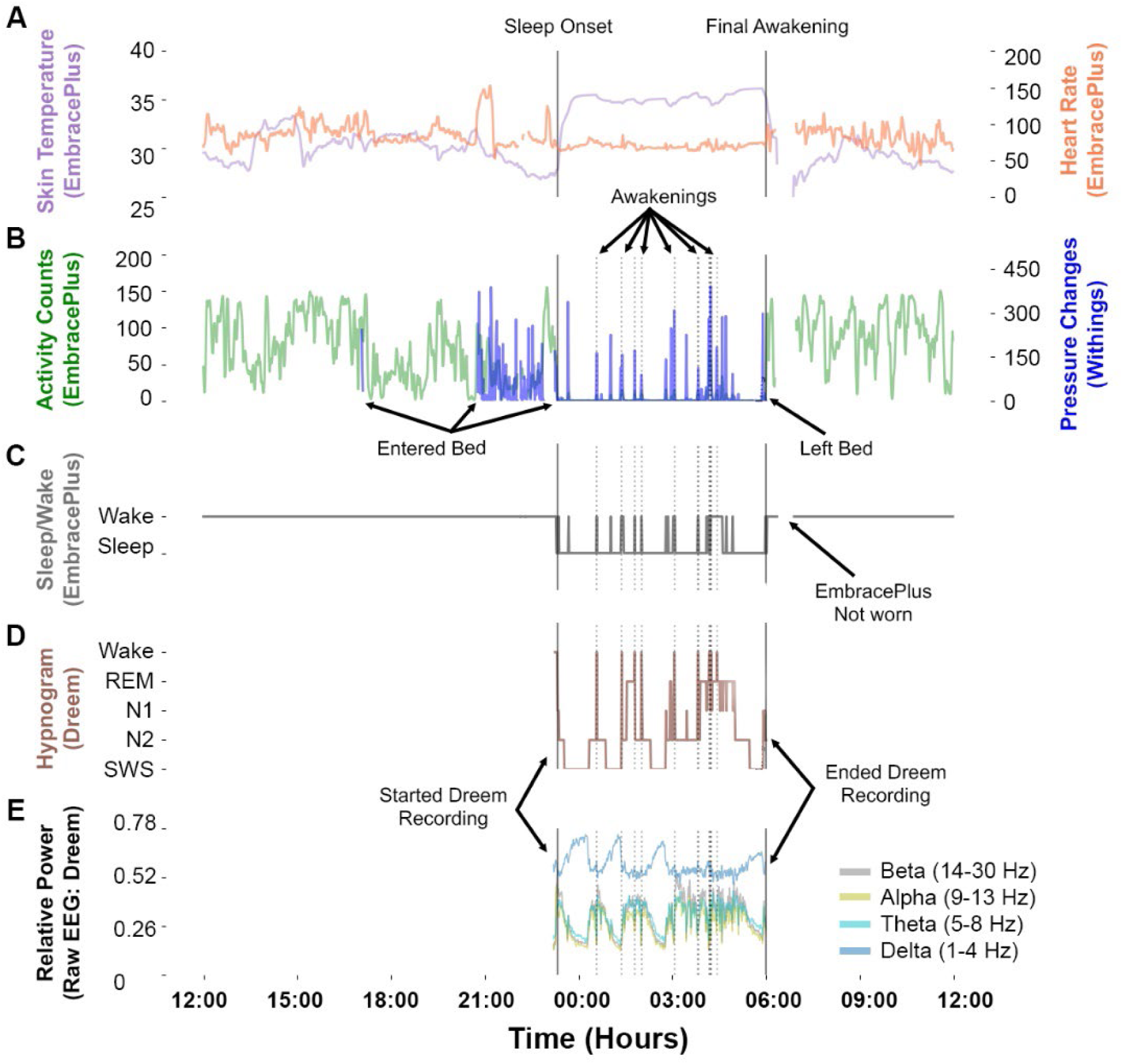
24-hour recording from a single participant starting and ending at noon. **A.** Skin temperature (purple) and heart rate (orange). **B.** Activity counts from EmbracePlus accelerometer (green) and pressure changes from the Withings Sleep mat (blue). **C.** Sleep/wake classification based on accelerometer data analyzed with EmbracePlus algorithms. **D.** Hypnogram from the Dreem3 automatic sleep staging algorithm. **E.** Average EEG power across all Dreem EEG channels. Average delta (1-4Hz, blue), theta (5-8Hz, turquoise), alpha (9-13Hz, yellow), and beta (14-30Hz, gray) band power demonstrate multiple sleep cycles throughout the night. Solid vertical lines mark sleep onset and final awakening as labeled. Dotted vertical lines mark awakenings during the sleep period according to the Dreem algorithm.

### Agreement across devices and with parent-reported sleep measures

We assessed agreement across devices in identifying four basic sleep measures (Figure 4): sleep onset (SO), final awakening (FA), wake after sleep onset (WASO), and total sleep time (TST, time from SO to FA minus WASO). We identified nights where all three devices recorded the child’s sleep successfully (see methods) and then extracted SO, FA, WASO, and TST as identified by the proprietary algorithm of each device per night. The same measures were also extracted from sleep diary data as reported by parents for the same nights. We limited our analyses to children with at least 3 nights of valid data from all devices and sleep diary (1151 nights from 69 autistic children and 67 siblings). This enabled us to also assess agreement between devices/diary measures that were averaged across at least 3 nights and parent-reported measures from questions in the CSHQ, where parents reported typical, average sleep measures per child.

**Figure 4:**
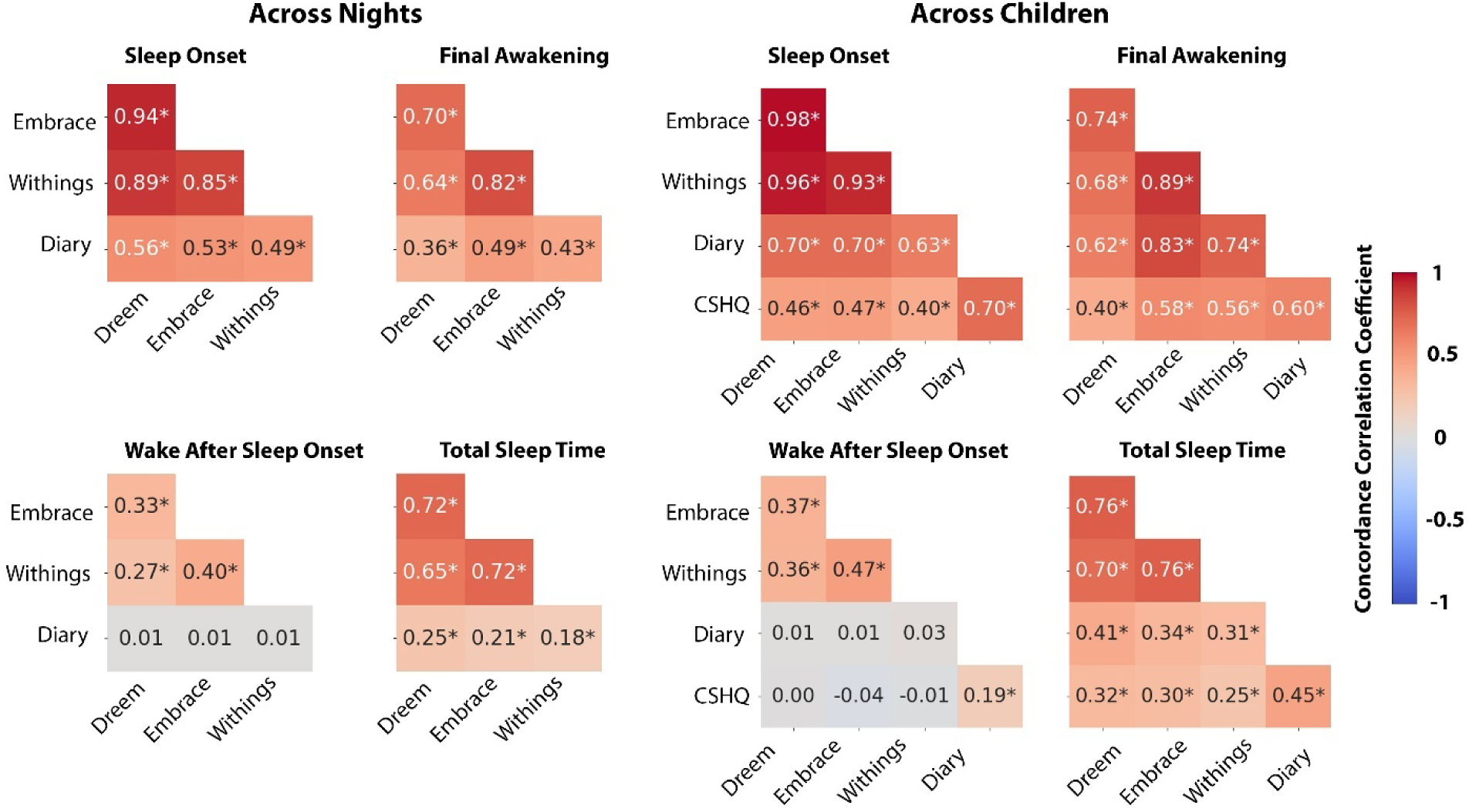
Concordance correlation coefficients (CCCs) demonstrating the pairwise agreement across devices, sleep diary, and CSHQ in estimating key sleep measures - Sleep Onset. Final Awakening, Wake After Sleep Onset, and Total Sleep Time. **A.** Computed across individual nights. **B.** Computed across participants after averaging measures across nights (per device/diary) and adding CSHQ data. CCCs are coded by color and values are noted for each pair. **Asterisks:** significant correlation (* p<0.001, randomization analysis)

We computed concordance correlation coefficients (CCC) to assess agreement between pairs of devices and/or sleep diary measures across nights (Figure 4A). This analysis demonstrated moderate to excellent agreement across the three devices when estimating SO, FA, and TST, with weaker yet still significant CCCs for WASO. There were considerably weaker, yet still significant CCCs between each of the devices and the sleep diary data for SO, FA, and TST, but not for WASO where there was no agreement between sleep diary and any of the devices.

Next, we performed the same analysis after averaging each measure across all available nights per child (Figure 4B). This yielded even stronger agreement across devices with moderate to excellent CCCs for SO, FA, and TST, as well as weaker yet still significant CCCs for WASO. Here too there were considerably weaker, yet still significant CCCs between each of the devices and sleep diary data for SO, FA, and TST, but not for WASO. Adding the CSHQ parent-reported measures revealed moderate CCCs with each of the devices for SO, FA, and TST, but not for WASO where there was no agreement. CCCs between CSHQ and sleep diary were significant across all measures including WASO with poor to moderate values.

These analyses demonstrated an important key finding: There was far stronger agreement across sleep measures from the three independent devices than there was between any of the devices and parent-reported measures from the sleep diary or CSHQ. This finding was particularly strong for WASO where there was no agreement between devices and parents regarding the extent of night-time awakenings of individual children whether examining individual nights or their mean per child.

### Comparison of device sleep measures across groups

To ensure consistency, we performed all further analyses with the same subset of data described above. We did not find any significant differences between groups in objective TST or WASO measures extracted from any of the three devices (Figure 5). TST differences between autism and sibling groups were, on average, <4.1 minutes according to all devices and were not significant (8.9>β>4.8, p > 0.23). WASO differences between groups were, on average, <3 minutes according to all devices and were also not significant (β<1.6, p > 0.66). Interestingly, ICC of TST values from all three devices were between 0.52-0.6, demonstrating that much of the variance in sleep duration was explained by differences across families rather than differences in diagnosis, age, or sex of the children. ICC ratios for WASO values were smaller with a range of 0.01-0.34.

**Figure 5:**
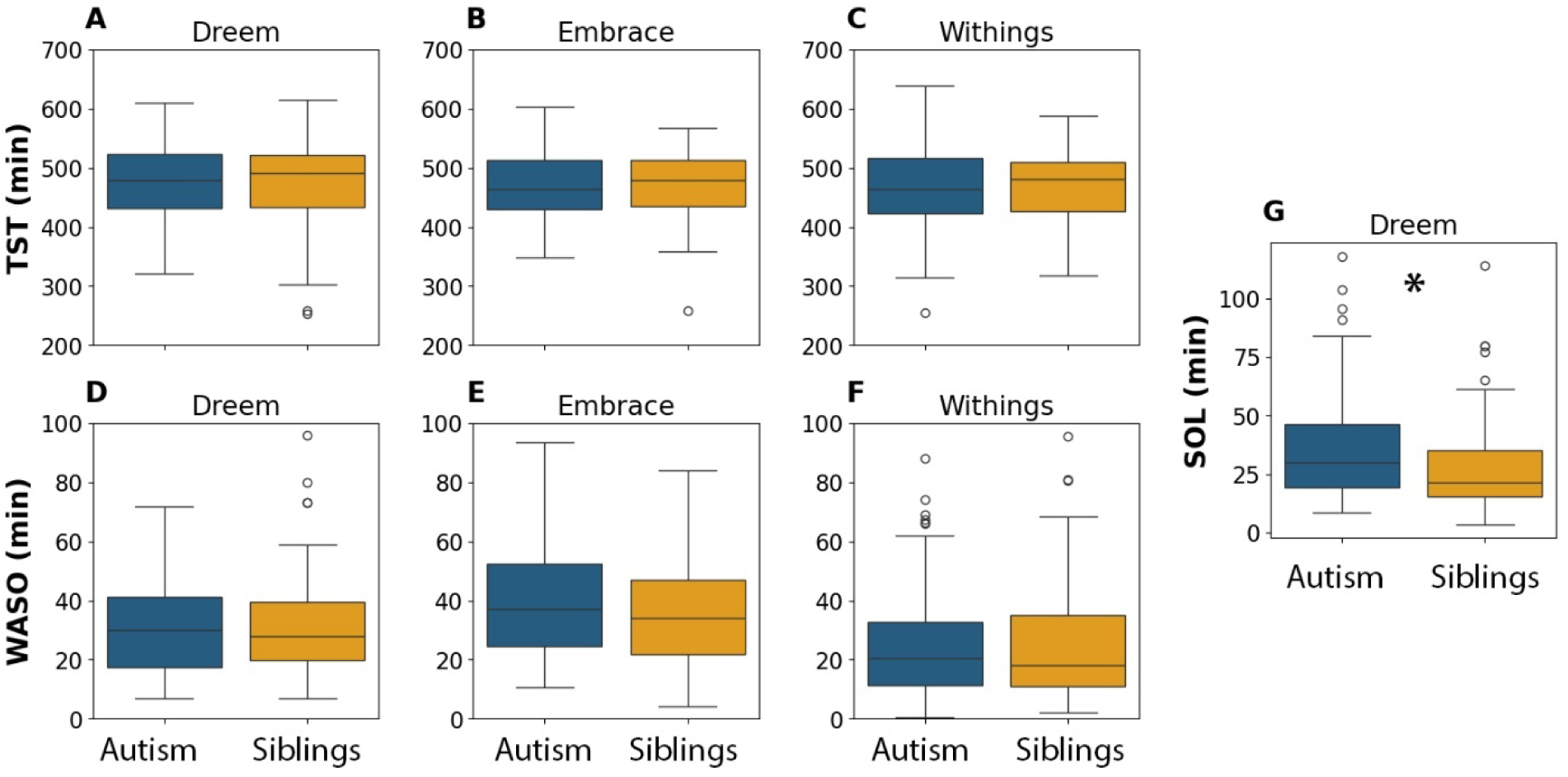
Comparison of total sleep time (TST), wake after sleep onset (WASO), and sleep onset latency (SOL) between autistic children (blue) and their siblings (yellow). **A.** TST from Dreem **B.** TST from Embrace **C.** TST from Withings **D.** WASO from Dreem **E.** WASO from Embrace **F.** WASO from Withings **G.** SOL from Dreem. **Asterisks:** significant differences between autism and sibling groups according to a mixed linear model analysis (p<0.05).

In contrast, sleep onset latency (SOL) differed significantly between groups. We defined SOL as the time from Dreem recording onset to sleep onset given that families were instructed to start the Dreem recording when the child was ready to go to sleep. We were unable to compute independent SOL measures from the EmbracePlus or Withings devices, because we did not have a clear indication of the time the child was ready to go to sleep from either of these devices. Dreem defined SOL was, on average, 8 minutes longer in autistic children relative to their siblings, a difference that was statistically significant (β = 7.56, p = 0.03). There were no significant effects of sex (β = -0.03, p = 0.99) or age (β = 0.11, p = 0.13) on SOL.

### Behavioral difficulties were associated with EEG derived SOL but not WASO

Given that EEG recordings enable direct identification of sleep/wake epochs and are used as the gold-standard for sleep research, we focused our final analysis on EEG estimates of SOL and WASO, which measure difficulties in sleep initiation and maintenance, respectively (Figure 6).

**Figure 6:**
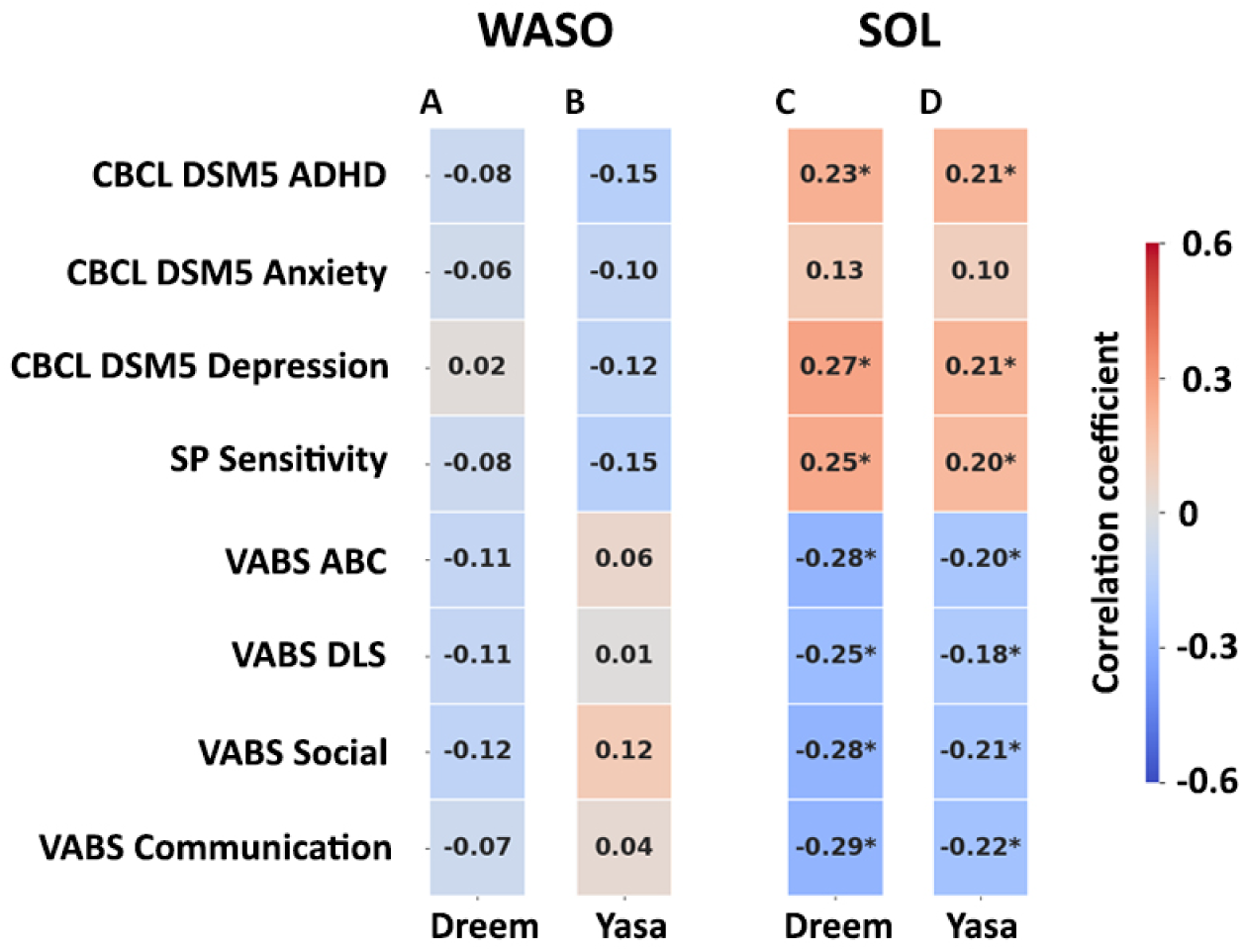
Matrix of Pearson correlation coefficients between EEG defined WASO and SOL measures (columns) and parent reported behavioral difficulties/abilities (rows). **A.** WASO according to Dreem algorithm. **B.** WASO according to Yasa algorithm. **C.** SOL according to Dreem Algorithm. **D.** SOL according to Yasa algorithm. Rows correspond to Child Behavior Checklist (CBCL), Sensory Profile (SP), and Vineland Adaptive Behavior Scale (VABS) subscale scores. **Asterisks:** significant Pearson’s correlation between the sleep and behavioral measures (p < 0.05, uncorrected).

We generated two independent estimates of each sleep measure with the Dreem and Yasa^39^ sleep staging algorithms, which yielded independent hypnograms per night/recording. Note that in this analysis we demonstrate the utility of the raw EEG data available in the SSP, which enabled us to validate the results of the Dreem sleep staging algorithm with Yasa, an open-source sleep staging algorithm^39^.

In this analysis we adopted a dimensional approach^40^ and examined whether sleep disturbances were associated with behavioral difficulties across study participants, regardless of autism diagnosis. We assessed the relationship between each sleep measure and multiple parent-reported behavioral measures, including CBCL subscales for DSM-5 ADHD, anxiety, and depression symptoms (three psychiatric symptom domains that are commonly associated with sleep problems), Sensory Profile subscale for sensory sensitivity (also commonly associated with sleep problems), and Vineland adaptive behavior scale scores that are indicative of general function.

Neither Dreem nor Yasa estimates of WASO were significantly correlated (-0.15<r<0.12, p>0.08) with any of the examined behavioral measures (Figure 6 A&B). In contrast, Dreem and Yasa estimates of SOL were significantly correlated with all behavioral measures (|r| > 0.18, p < 0.04) except for CBCL anxiety scores (r < 0.13, p > 0.13). These results demonstrate a reliable dissociation between EEG-defined WASO and SOL, suggesting that sleep initiation problems, rather than sleep maintenance problems, are significantly associated with a variety of behavioral difficulties and poorer adaptive behaviors, regardless of autism diagnosis.

## Discussion

This manuscript presents the extensive data available in the SSP, which includes raw and processed, high-resolution, synched data from multiple devices/sensors (Figures 1 & 3) along with parent-report questionnaires and daily sleep diaries (Figure 2). These data offer rich, unique opportunities to study sleep behavior and neurophysiology as well as daytime behaviors in autism. The data also offer broad opportunities for methodological development including training deep learning algorithms to identify and quantify a variety of behaviors and states (e.g., hyperactivity). Furthermore, whole exome sequencing data is available for all participants and their parents, enabling future investigations of genotype-phenotype relationships.

An important feature of the SSP is the simultaneous collection of multiple objective and subjective measures of sleep that enable their comparison. Initial analyses of common sleep measures including SO, FA, WASO, and TST demonstrated considerably larger agreement across device pairs than between each device and parent-reported sleep diary or CSHQ measures (Figure 4). Averaging measures across multiple nights per participant increased agreement across devices, demonstrating the value of multi-night recordings for reducing device measurement errors on individual nights. Most importantly, these findings reveal that objective sleep measures, acquired with three independent devices and sensor types, yielded more accurate and reliable data than subjective parent reports. Note that device recordings included sleep EEG data, which is typically used as “ground truth” for classifying sleep/wake periods. This motivates further research with multi-device, multi-night study designs that can accurately quantify sleep in the home environment.

Parents reported greater sleep problems in autistic children relative to their siblings, including significantly larger WASO and higher total sleep disturbance scores (Figure 2). However, when comparing objective sleep measures from the devices, we found that only SOL was significantly longer in the autism group (Figure 5). Given the limited agreement between device and parent-reported sleep measures (Figure 4), this suggests that parent-reported sleep disturbances such as prolonged WASO are not necessarily evident in direct, objective measures of sleep. Indeed, a recent meta-analysis of actigraphy studies in autism also reported that SOL was significantly longer in autistic children relative to typically developing controls while WASO was not^2^. PSG studies have also mostly reported no difference in WASO across groups^21–24^.

Given that siblings of autistic children are known to have elevated rates of developmental and behavioral difficulties relative to the general population (see following section), we also analyzed the data using a dimensional approach^37,40^. Here, we examined whether the severity of common co-occurring psychiatric symptoms (ADHD, anxiety, and depression) as estimated by the CBCL, sensory sensitivity as estimated by the Sensory Profile, and adaptive function as estimated by the Vineland, were related to two key sleep disturbance measures: WASO and SOL. We performed this analysis independently with two sleep staging algorithms (Dreem and Yasa) and demonstrated equivalent results across both. While SOL was significantly correlated with multiple behavioral measures, WASO was not (Figure 6). This dissociation highlights the need to further study sleep initiation problems and their underlying physiological mechanisms in children with behavioral and developmental difficulties, while using direct, objective sleep measures.

The examples above demonstrate a small subset of potential sleep analyses that can be explored using processed or raw data in the SSP. Further studies could examine additional sleep measures including sleep regularity (i.e., across nights), circadian rhythm, sleep architecture, and sleep neurophysiology (e.g., sleep spindles) that may hold important insights for basic and clinical autism research.

### The SSP sibling study design

The sibling study design implemented in the SSP is both a feature and a limitation of this resource. Parents reported that participating autistic children exhibited significantly greater behavioral difficulties than their non-autistic siblings with particularly large differences in the severity of core autism symptoms, as expected (Figure 2). Nevertheless, siblings of autistic children are known to exhibit elevated rates of developmental and behavioral problems than the general population, including high rates of ADHD, language delays, cognitive difficulties, and mental health problems^34,35^. Indeed, parents of children in the SSP reported that 58% of autistic children and 30% of siblings had a medical diagnosis of ADHD (Table 1). Moreover, multiple studies have reported that siblings often have mild autism symptoms that place some of them within the broad autism phenotype^41^.

When studying autism using SSP data it is, therefore, important to consider that sibling data is not equivalent to “control” data from children free of any developmental concerns, as is common in categorical case-control studies. Lack of significant differences in TST between autism and sibling groups (Figure 2) should not be interpreted as evidence for lack of sleep disturbances in autistic adolescents. Instead, both autism and sibling groups may exhibit reduced TST in comparison to children without developmental concerns due to high rates of other psychiatric conditions (e.g., ADHD) which are also associated with sleep disturbances^4^. The SSP may therefore be particularly useful for studies that adopt a dimensional or transdiagnostic approach with the goal of unraveling relationships between multiple behavioral and neurophysiological dimensions regardless of categorical diagnoses (e.g., Figure 6).

While the sibling design may seem to introduce some complexity, it simultaneously affords several advantages. First, the sibling design constrains multiple environmental and familial factors that are usually not accounted for in case-control study designs. For example, sleep behavior is strongly influenced by familial genetics^42^ and sleep hygiene^43^. Such factors are likely to differ across autism and unrelated control groups yet are constrained in autism-sibling pairs. Indeed, our mixed-model analyses of all sleep measures included sibling pairs as a random-effects factor and revealed that >50% of the variance in TST values (in all 3 devices) was explained by between-family differences rather than autism-sibling differences. A second advantage is that subjective parent reports were collected for both diagnostic groups from the same parents, thereby reducing between-group differences in reporter bias^44^. These advantages suggest that when significant differences across autism-sibling pairs are detected in this cohort (e.g., SOL, Figure 5), they are likely to be of high validity given that multiple environmental factors are controlled. A final additional advantage is the availability of whole exome DNA sequence data from all participants and their parents, offering the opportunity to use familial genetic techniques to test for potential gene-phenotype relationships, although such research will likely require expanding the number of participants in the SSP to enable higher statistical power.

### The remote at home SSP study design

Previous autism sleep studies have mostly used questionnaires, actigraphy devices, or PSG recordings that were performed during a single night, mostly in a sleep lab. These past studies typically focused on one measurement technique at a time with actigraphy and PSG studies often further limited by small samples^2,21–25^.

The SSP combines multiple research techniques in a remote study design where wearable devices were mailed to the families who operated them independently at home. This design had several key advantages over previous studies. First, data were collected at home (i.e., high ecological validity) rather than in a sleep lab. This is particularly meaningful in autism research, because autistic participants often find changes in routines particularly difficult^28^ such that sleep measurements from a sleep laboratory may not represent typical sleep at home. Second, because data were collected with a wearable EEG device rather than a traditional PSG system, we were able to record most participants across multiple days/nights. This was important for ensuring that autistic participants acclimated to the EEG devices^45^ and critical for allowing future studies to calculate measures of sleep regularity and circadian rhythm that require multiple nights of data. Third, since the SSP utilized multiple devices in parallel it is possible to compare data (e.g., Figure 4) and demonstrate the reliability of findings across devices, addressing important concerns regarding reproducibility^46^. Fourth, remote data collection enabled us to record data from up to thirty families in parallel; performing an equivalent study in a sleep laboratory would have required significantly greater resources. Fifth, all our devices continuously streamed data to the device company servers, enabling us to monitor data acquisition and troubleshoot with families when necessary. This is the first remote sleep study in autism to apply such techniques for ensuring high quality, continuous data collection from multiple devices. Finally, broadly speaking, remote data collection can afford families in isolated locations and those in low socioeconomic situations an equal opportunity to participate in research. The current study included families from 22 states throughout the U.S., including those living in remote rural areas.

Despite the many advantages listed above, remote studies also introduce a variety of challenges. First, some participants were not able to successfully record multiple nights with all devices. Hence, only a subset of the collected data was used when comparing data across devices (Figures 4 and 5). Note that future studies examining individual device data or even pairs of devices will be able to work with considerably larger subsets of the data than included in our stringent analyses. Second, because the selected devices are relatively new and have been validated in only a relatively small number of studies^32,47,48^, their measures may not be directly comparable to those of older actigraphy devices or lab-based PSG systems. Indeed, the Dreem EEG data includes only a partial PSG montage with no EOG, EMG, or mastoid channels, thereby limiting the analyses that can be performed with this data. Third, devices that can be used remotely are often suitable only for older children and adults. For example, the Dreem headband requires a minimal head circumference of 52cm and is not likely to fit most children under the age of 10. Hence, some devices implemented in the SSP will not be appropriate for remote studies of younger cohorts. Finally, remote data collection is far less controlled than research in the lab. Participants may inadvertently rotate or remove devices in the middle of the night or use them in unexpected ways when they are not monitored by researchers. Hence it is important to assess data quality and define clear criteria for exclusion. Initial steps to enforce such criteria in the SSP are exclusion of recordings where TST was less than 3 hours or exceeded 16 hours, WASO exceeded 3 hours, and/or recordings where EEG was of low quality. Hence, an important principle of remote data collection is to acquire extensive data with the expectation that it will contain anomalies and noise that will require careful cleaning and harsh exclusion criteria. Ideally, researchers will formulate open-source processing pipelines for SSP data that can be shared so that additional data cleaning steps are standardized across future studies.

### Digital phenotyping with wearable/nearable devices: Beyond sleep

In the initial analyses of the SSP, we focused on objective quantification of sleep disturbances in autism, a commonly reported co-occurring condition of high priority to autistic individuals and their families. However, the SSP offers a wide variety of data for studying a variety of behaviors beyond sleep. For example, recent studies have utilized electrodermal activity (EDA) recordings to identify tantrums and emotional outbursts in autistic children with challenging behaviors^5^ while others have used actigraphy to quantify hyperactivity in children with ADHD^49^. The availability of raw accelerometer, EDA, skin temperature, and photoplethysmography (PPG) data from multi-day EmbracePlus recordings along with daily parent-reported data on tantrums and emotional regulation may enable a variety of studies on these topics in autism using the SSP dataset.

### Conclusions

The SSP is a new open-science resource that affords researchers from multiple disciplines a unique opportunity to study human behavior, sleep, and autism using digital phenotyping techniques. It demonstrates the feasibility of recording sleep and daily behavior remotely from a relatively large sample of adolescent participants with and without autism, using multiple synchronized devices in parallel over an extended period. Initial analyses demonstrate the power of this dataset and reveal the specific prominence of sleep initiation problems in autism as well as their association with a variety of developmental and behavioral problems in both autistic children and their siblings. SSP data, analysis code from this study, and whole-exome sequencing data from all participants and their parents are available through SFARI base. We encourage the research community to take advantage of these free, open resources and hope the SSP will be used widely to advance research in autism and beyond.

## Online Methods

### Participants

All participating families were recruited from the Simons Powering Autism Research (SPARK) cohort^27^, which includes over 50,000 individuals with autism in the U.S., many of whom have contributed DNA samples for whole exome sequencing and completed basic medical history and other questionnaires. In many cases, immediate family members have also joined SPARK, contributed DNA samples, and consented to be re-contacted for participation in additional studies. SPARK families were contacted by email with the opportunity to join the SSP if they had two adolescent children who were full siblings, 10-17 years old, one with a parent-reported medical diagnosis of autism and the other without. In addition, all children had completed whole-exome sequencing, which did not yield any returnable autism-related genetic results (i.e., all autistic participants had idiopathic autism) and all autistic children exceeded the autism cutoff score (>15) on the Social Communication Questionnaire, Lifetime Edition, which was completed by a caregiver at the time they joined SPARK. All parents completed informed consent, and the study was approved by the WCG IRB committee.

A total of 2609 SPARK families fulfilling the inclusion criteria described above were contacted by email and 315 families completed the informed consent and were recruited. Of these, 113 families completed all baseline questionnaires for both children and were sent wearable devices. Eleven of these families dropped out of the study for different reasons including difficulties using the devices, familial difficulties, and change of heart. A total of 102 families, living in 22 states throughout the U.S. successfully completed the study with data recorded from 102 autistic children and 98 siblings (Table 1).

### Procedure

Recruited families completed nine parent questionnaires for each of their children before receiving the devices (Table 1). These included the Child Sleep Habits Questionnaire (CSHQ)^28^, Vineland Adaptive Behavior Scales^29^, Child Behavior Checklist (CBCL)^30^, Aberrant Behaviors Checklist (ABC)^31^, Social Responsiveness Scale, 2^nd^ edition (SRS-2)^32^, Repetitive Behaviors Scale Revised (RBS-R)^33^, Sensory Profile, 2^nd^ edition^34^, Family Inventory of Sleep Habits (FISH)^35^, and a short medical update questionnaire where parents were asked to report on medically diagnosed conditions (beyond autism) for each of their children (Supplementary Table 1).

Families were then sent a package with the Dreem3 EEG headband (*Beacon Inc.*), EmbracePlus smartwatch (Empatica Inc.), Withings Sleep mat (*Withings Inc.*), a Samsung A51 smartphone set up with apps for the three devices to enable continuous data transfer, and written instructions on how to use all devices. Study coordinators completed an onboarding procedure with each family, ensuring that data collection began with the autistic child and instructing one of the parents to complete a daily sleep diary online, using a link from a daily text message reminder. Parents also reported on medications taken during the study as part of the daily sleep diary (Supplementary Table 2). We grouped medications according to their clinical indication (Supplementary Table3) when presenting results across autism and sibling groups (Table 1).

Data collection was monitored remotely, and families were contacted when data was missing or of low quality. After approximately 3 weeks of data collection from the autistic child, devices were transferred to their sibling and data collection continued for another 3 weeks. We prioritized data collection from the autistic participants who were always recorded first, because this data was more valuable for the purposes of the study. Finally, the devices were returned by courier, and the families were given a gift card and sent a sleep report summarizing some of the data collected from each of their children.

### Data acquisition and structure

The Dreem3 headband recorded EEG data from 5 channels over frontal and occipital cortex (F1, Fz, F2, O1, and O2) at 250Hz as well as accelerometer data at 50 Hz. Participants were instructed to turn on and wear the device before going to bed and turn it off when they woke up in the morning. Data was uploaded automatically to Beacon Inc. servers every time the headband was charged. Their automated system then applied a bandpass filter with a low cutoff frequency of 0.4Hz and a high cutoff frequency of 35Hz, generating a single H5 and EDF file with the raw EEG per night. In addition, the Beacon system also automatically generated a hypnogram file per night in txt format using their proprietary sleep staging algorithm and a summary statistics file with common sleep measures such as TST and WASO per night. Recordings containing less than 3 hours or more than 16 hours of TST were discarded from the SSP and are not reported in this manuscript. Further details regarding Dreem3 data acquisition and structure are available on their website (https://beacon.bio/dreem-headband).

The EmbracePlus smartwatch recorded data from 4 sensors: accelerometer (at 64Hz), photoplethysmography (PPG, at 64Hz), electrodermal activity (EDA, at 4Hz), and skin temperature (at 1Hz). Raw data were continuously transmitted from the device to the Care Lab app on the cellphone via Bluetooth and then automatically transferred and stored on the Empatica servers in JSON files of variable lengths. We concatenated JSON files with raw data from each sensor into a single CSV file per 24 hours per sensor starting at 12am according to their timestamps. Segments without data (e.g., participants took off the watch) were replaced with not-a-number (NaN) values to maintain equivalent file structure. In addition, Empatica automatically computes summary statistics of the data which include, for example, step-count, heart rate (HR), HR variability, and sleep-wake classification per minute. These data were also reformatted into CSV files with the same 24-hour structure as the raw data. Further details regarding EmbracePlus data acquisition and structure are available on their website (https://www.empatica.com/embraceplus).

The Withings Sleep mat contains an inflated chamber that is placed under the mattress in the area corresponding to the participant’s torso. Raw data are not available from the device, but pressure changes, HR, movement statistics, and sleep-wake classification measures are available per minute (0.016 Hz). These data are recorded only when the participant is in bed and stored in a separate CSV file per night. Recordings with less than 3 hours or more than 16 hours of TST were discarded from the SSP and are not reported. Further details regarding Withings data acquisition and structure are available on their website (https://www.withings.com/us/en/sleep).

Finally, a designated parent received SMS text messages every morning and evening with 6-8 questions about the child’s behavior during the preceding night and day, respectively as well as a question regarding medication intake (Supplementary Tables 2&3). Sleep diary was successfully collected for 188 of the participants (ASD = 95, Siblings=93).

### Data harmonization

Harmonization of data across the three devices and sleep diary involved several steps. First, collected data, initially organized by device user ids, was mapped to SPARK participant IDs. Second, data time stamps were adjusted to the participants’ local time zone. Third, nightly Dreem and Withings recordings were labeled according to the date of final awakening and recordings with less than 3 hours of TST or more than 16 hours of TST were excluded. Fourth, EmbracePlus data were concatenated and split into separate files containing 24 hours of data. Files with less than 6 hours of data were excluded. All data were transformed into equivalent CSV format, except for raw EEG data that was kept in EDF format. Additional information about filenames, data format, and data structure are available when accessing the data repository.

### Data Analysis

Initial analysis of collected data reflects the total number of recordings/files available from each device and their overlap on the same dates (Figure 1). Similarly, initial analysis of questionnaire data and comparison across autism and sibling groups included all participants in the SSP (Figure 2). To demonstrate the data structures described above we arbitrarily selected a participant with a 24-hour period that contained data from all three devices and displayed selected measures from the three devices according to the participants’ time (Figure 3).

All further analyses focused exclusively on data from nights where data were available from all three devices and sleep diary. These analyses included data from 69 autistic children and 67 siblings who successfully recorded at least 3 valid nights with all devices and sleep diary. A valid night recording included a TST of 3 to 16 hours that started between 7pm and 7am and WASO that did not exceed 3 hours from each of the devices. All nights that did not fulfill these criteria were excluded with the assumption that they contained extreme values that were likely caused by device failures (e.g., running out of power) or algorithm errors. In addition, Dreem recordings with a quality score of ≤30 out of 100 (as computed by Beacon) were also removed. Using these criteria, we isolated 1151 nights with high-quality simultaneous data recordings from all three devices and sleep diary.

Dreem and Withings algorithms yield estimates of SO, FA, WASO, and TST as part of their nightly reports. EmbracePlus algorithms, however, only yield sleep/wake labels per one-minute epoch of recording, rather than overall nightly reports. We calculated SO, FA, WASO, and TST from the EmbracePlus data using the following criteria. SO was defined as the first sleep epoch followed by at least 10-minutes of consistent sleep. FA was defined as the first wake epoch after the last 10-minute segment of sleep that was followed by at least two hours of wake. WASO was calculated as the sum of wake epochs between SO and FA that were in wake segments that were 3 minutes or longer (i.e., at least 3 minutes long). TST was calculated as the time between SO and FA minus WASO. SOL was computed for Dreem data as the time between recording start and SO.

Parents reported SO, FA, and WASO per night as part of the sleep diary, enabling us to compute daily TST (FA-SO-WASO). Parents also reported SO, FA, and WASO on the CSHQ, but while SO and FA were reported separately for weekdays and weekends, WASO was reported regardless of day type. We, therefore, computed the mean SO/FA across weekdays and weekend before computing TST (FA-SO-WASO).

In our final analysis, we also extracted independent SO, FA, WASO, and TST values from hypnograms that were computed from the raw EEG recordings using the Yasa sleep staging algorithm^39^. We applied the Yasa algorithm to raw EEG data from all available channels in each EDF file (i.e., night). Yasa yields a sleep stage score per 30-second epoch per channel indicating its confidence in identifying the proper sleep stage within a given epoch. We identified a single sleep stage per epoch by computing the majority vote across all channels, thereby yielding a hypnogram per night. SO was defined as the first epoch in the first 10-minute segment of sleep. FA was defined as the first wake epoch following the last 10-minute segment of sleep that was followed by end of recording or two hours of wake. TST was calculated as the time between SO and FA, excluding WASO. WASO was calculated as the sum of wake epochs between SO and FA.

### Data Sharing

All SSP data is available through SFARI Base at https://base.sfari.org/dataset/DS0000089. All processed data and code used to generate the results and figures presented in the current manuscript are available at: https://github.com/Dinstein-Lab/SSP_manuscript

### Statistical Analysis

All statistical analyses were performed using custom written Python code. Comparison of questionnaire scores and sleep measures across groups were performed with mixed linear model analyses as implemented in the Statsmodel Python package. We used family ID as a random effect and diagnosis (autism or sibling), age, and sex as fixed effects. This enabled us to separate potential differences across families that are likely to affect both siblings living in the same household from individual effects of diagnosis, age, and sex. We also computed Intraclass Correlation Coefficients (i.e., between family variance divided by the sum of between and within family variance) to determine whether differences between families were smaller or larger than differences among individuals.

We computed concordance correlation coefficients (CCCs) to assess pair-wise agreement across sleep measures from the different devices, sleep diary, and sleep questionnaire (CSHQ). CCC is a common measure of reproducibility where error is calculated as distance from the diagonal (absolute agreement) rather than from a linear fit line (relative agreement). To determine the significance of CCCs we performed a randomization analysis where we created a distribution of CCCs expected by chance when shuffling values across nights (i.e., shuffling date labels) in 5000 different iterations. For a CCC to be considered significant it had to fall beyond the 0.1 top percentile of this distribution (equivalent to a p value of 0.001). We chose this conservative p value to correct for the 10 multiple comparisons, across devices, sleep diary, and CSHQ, performed for each sleep measure.

Finally, we performed Pearson correlation analyses between questionnaire scores and WASO or SOL measures as extracted from Dreem recordings with the Dreem or Yasa algorithms. Statistical significance was set to a p value of 0.05 and we did not correct for multiple comparisons in this analysis to maintain sensitivity. Note that the goal of this analysis was to determine whether consistent relationships with behavior were apparent for either sleep measure across the two sleep staging algorithms.

## Acknowledgements

This study was funded by the Simons Foundation Autism Research Initiative (SFARI). Thanks to the following advisors who helped shape the Simons Sleep Project: Maja Bucan, Dara Manoach, Mathew Goodwin, Dirk-Jan Dijk, Ashura Buckley, Ullrich Bartsch, Max de Zambotti, Michael Snyder, Edward Brodkin, David Nobbs, Emily Simonoff, Guillermo Sapiro, and Jonathan Sebat. Thanks to the SPARK and SFARI informatics teams for supporting the project and to Kelsey Martin, Sachin Ranade, Jennifer Foss-Feig, Aaron Wong, and Swami Gansen for constructive conversations about the design and implementation of this project.

