## Supplemental Materials for "Simons Sleep Project (SSP): An open science resource for accelerating scalable digital health research in autism and other psychiatric conditions"

* Corresponding author

**Supplementary Table 1:** Medical update questionnaire

| |
|---|
| **1. Has your child been diagnosed with any of the following sleep disorders by a physician?** (Please select all that apply) |
| • None |
| • Insomnia |
| • Sleep Apnea |
| • Sleep Arousal |
| • Nightmare Disorder |
| • Sleep Behavior Disorder |
| • Hypersomnolence Disorder |
| • Narcolepsy |
| • Restless Legs Syndrome |
| • Circadian Rhythm Sleep Wake Disorders |
| **2. Has your child been diagnosed with any of the following behavioral disorders by a physician?** (Please select all that apply) |
| • None |
| • ADHD/ADD |
| • Conduct Disorder |
| • Intermittent Explosive Disorder |
| • Oppositional Defiant Disorder |
| • Repeating Bowel Accidents |
| **3. Has your child been diagnosed with any of the following neurological disorders by a physician?** (Please select all that apply) |
| • None |
| • Brain Infection |
| • Seizures Epilepsy |
| • Traumatic Brain Injury |
| • Tourette Syndrome |
| **4. Has your child been diagnosed with any of the following psychiatric disorders by a physician?** (Please select all that apply) |
| • None |
| • Eating Disorder |
| • Alcohol or Substance Use |
| • Personality Disorder |

| |
|---|
| • Anxiety Disorder |
| • Bipolar Disorder |
| • Depression |
| • Dysregulation Disorder |
| • Hoarding |
| • Obsessive Compulsive Disorder |

**Supplementary Table 2:** Sleep diary questions

| Section | Question | Response options |
|---|---|---|
| **Evening Diary (PM)** | How energetic was your child today? | **Scale 0-10** (Very tired to Highly energetic) |
| | How was your child's mood today? | **Scale 0-10** (Terrible to Great) |
| | How did your child feel physically today? | **1**=Great, **2**=Average, **3**=Minor aches/pains, **4**=Didn't feel well, **5**=Was sick |
| | Did they have a fever? | **Yes/No** |
| | Any Gastro-intestinal problems? | **Yes/No** |
| | Did your child have any tantrums today? | **Yes/No** |
| | **If yes:** How many tantrums? | **Number** (1-10) |
| | My child took these medications today | **Free text:** medication name and dosage |
| | My child consumed caffeinated items in the: | **Multiple choice**: Morning, Afternoon, Evening, None |
| | My child exercised for at least 20 minutes in the: | **Multiple choice**: Morning, Afternoon, Evening, None |
| | My child took a nap today? | **Yes/No** |
| | **If yes:** Nap time | **Time of day** Hour:Minute (AM/PM) |
| | **If yes:** Nap duration | **Time** Drop-down (15-minute intervals) |
| **Morning Questions (AM)** | My child got into bed last night at: | **Time of day** Hour:Minute (AM/PM) |
| | How anxious/restless was your child at bedtime? | **Scale 0-10** (calm/relaxed to anxious/restless) |
| | I noticed that my child woke up during the night (X) times | Numerical value |
| | Total minutes awake during night | **Time** Numerical value (Minutes) |
| | When your child woke up during the night, they: | **Multiple choice**: Left bed, Awakened parents, Required help, Not applicable |
| | What was the reason for waking up? | **Multiple choice**: Noise, Light, Bad dream, Bathroom, Agitated/anxious/emotional, Not applicable |
| | This morning my child woke up at: | **Time of day** Hour:Minute (AM/PM) |
| | When my child woke up for the day, he/she felt: | **Scale 0-10** (Rested to Tired) |

**Supplementary Table 3:** Categorization of parent-reported medication brand names to medication groups (Table 1)

| Category | Medication Brand Name | | |
|---|---|---|---|
| **Antidepressants** | Prozac (fluoxetine)<br>Zoloft (sertraline)<br>Lexapro (escitalopram) | Trazodone<br>Cymbalta (duloxetine)<br>Wellbutrin (bupropion) | Amitriptyline<br>Venlafaxine<br>Imipramine |
| **Stimulants** | Adderall<br>Vyvanse<br>Focalin (dexmethylphenidate)<br>Ritalin (methylphenidate) | Dextroamphetamine<br>Jornay<br>Concerta | Quillivant<br>Azstarys<br>Lisdexamfetamine |
| **Anti-anxiety** | Hydroxyzine<br>Buspar (buspirone)<br>Ativan (lorazepam)<br>Clonidine | Strattera (atomoxetine)<br>Depakote<br>Guanfacine<br>Intuniv | Oxcarbazepine<br>Lamictal (lamotrigine)<br>Tegretol (carbamazepine) |
| **Anticonvulsants** | Trileptal<br>Depakote | Lamotrigine<br>Carbamazepine | Oxcarbazepine |
| **Antipsychotics** | Abilify (aripiprazole)<br>Risperidone (Risperdal) | Seroquel (quetiapine)<br>Olanzapine (Zyprexa) | Vraylar<br>Ziprasidone |
| **Sleep Aids** | Melatonin | | |
